# TRPA1 channel activation by synthetic lipid nanoparticles

**DOI:** 10.64898/2026.05.03.722497

**Authors:** Alina Milici, Justyna B. Startek, Geert Bultynck, Karel Talavera

**Affiliations:** Laboratory of Ion Channel Research Dept. of Cellular and Molecular Medicine, KU Leuven, Belgium; Laboratory of Molecular and Cellular Signaling, Dept. of Cellular and Molecular Medicine, KU Leuven, Belgium

## Abstract

TRPA1 is a polymodal ion channel receptor known for its role in nociception. TRPA1 can be activated by local mechanical perturbations in the surrounding plasma membrane (PM) by molecules that insert in the lipid bilayer. Here, we tested whether TRPA1 function can be modulated by lipid nanoparticles (LNPs) while interacting with the target cell plasma membrane. We found that LNP induce irregular Ca^2+^ transients in heterologous and native TRPA1-expressing cells, which may reflect stochastic LNP-PM interactions. By using different cell types and applying selective and non-selective TRPA1 inhibitors, we revealed that the cytosolic [Ca^2+^] is elevated transients arise as a result through multiple mechanisms: TRPA1-dependent Ca^2+^ influx, TRPA1-independent Ca^2+^ influx, and Ca^2+^ mobilization from the endoplasmic reticulum. Our results describe a novel, non-canonical TRPA1 activation mechanism by LNPs, that may be relevant in the context of the development of cancer and nasal vaccines.

## Introduction

The proper function of organisms relies on their ability to react to changing environmental conditions, which critically depends on the ability to detect external stimuli. One example of molecular sensors that contribute to this process is the superfamily of Transient Receptor Potential (TRP) channels. TRP channels are localized in the plasma membrane (PM) of cells and act as sensors for chemical, thermal and mechanical cues [1–5]. TRPA1, the only mammalian member of the ankyrin subfamily, is arguably the most versatile polymodal sensor and is mainly localized in primary sensory neurons, where it is known for its major role in nociception [6,7].

TRP channels, and in particular TRPA1, can be modulated by the lipid composition of the PM and activated by mechanical membrane perturbations induced by bacterial lipopolysaccharides (LPS) and other lipophilic compounds [4,8–11]. Interestingly, rather than the classical compound-channel binding, LPS seem to activate TRPA1 by altering the organization of the surrounding plasma membrane by inserting between phospholipids, decreasing membrane fluidity and lateral compression of the channel [11]. A similar activation pattern was described in the case of alkylphenols [10]. Their effects on TRPA1 are proportional to the lipophilicity and related to local perturbations induced in the PM, rather than covalent binding to the channel. In the membrane, TRPA1 interacts with cholesterol in specialized structures called lipid rafts, and lipid raft destabilization alters the sensitivity of the channel to several agonists [12–14]. Based on the evidence presented above, the activity of TRPA1 is modulated by the local mechanical properties of the plasma membrane, either via its composition (i.e. the amount of cholesterol in lipid rafts), or via mechanical alterations of the plasma membrane by the insertion of lipophilic chemical moieties. This highlights the role of this channel as cellular sentinel operating as broadly-tuned sensors of mechanical perturbations of the plasma membrane [15].

Recent evidence shows that TRP channels can also be modulated by particulate matter [16]. This raises the possibility to be activated by cellular events involving membrane perturbations, such as those induced by synthetic lipid particles (e.g. ionizable lipid nanoparticles (LNPs)). LNPs are engineered lipid structures widely used as drug and vaccine delivery systems [17,18]. In the recent years, lipid assemblies forming lipoplexes with nucleic acids or small molecule drugs have gained tremendous popularity in biotechnology due to their advantages over classic delivery systems [19], and are now being deployed in multiple pre-clinical and clinical stages of gene- and small molecule-based therapeutics [18]. Although still early stages of development, efforts are being made to use the ionizable LNP technology for applications such as cancer and nasal vaccines [20,21]. A common hallmark of the tumor environment, as well as the upper airway mucosa is the relatively low pH (tissue acidosis in the case of tumors and the protective, slightly acidic environment of the upper airways) [22,23]. Under these conditions, LNPs formulated with ionizable lipids will be protonated in contact with the tissues, increasing the likelihood of interaction with the plasma membrane of target cells. Common to these two scenarios is the expression and functional implication of TRPA1. There is evidence that TRPA1 is upregulated or activated in different types of tumors, where it plays different roles in the tumor progression and therapeutic outcome, as well as cancer-induced pain [24–26]. Moreover, airway sensory nerves express a myriad of sensory receptors, including TRPA1 [27–30].

Throughout the LNP uptake process, LNP interactions with the PM cause mechanical perturbations via processes such as surface adsorption, endocytosis or fusion [31]. The long-term effects of LNPs on target cells have been thoroughly investigated. However, most studies have focused on cell and tissue functions (e.g. lysosomal function, lipid metabolism, release of factors in the extracellular medium, gene regulation, chronic inflammation, and immune activation and clearance) [32–36]. In contrast, the efforts to understand the acute effects (i.e. within several minutes) of lipoplexes on membrane sensory proteins remain limited, and almost exclusively related to the activation of Toll-Like Receptor 4 (TLR4) [37]. To our knowledge, there are yet no studies examining the acute effects of LNPs on TRP channels. However, two papers have described acute TRP channel modulation by endogenously released lipid particles, i.e. extracellular vesicles (EVs). According to these studies, interactions between EVs and target cells trigger intracellular Ca^2+^ signals via TRPM7 in microvascular endothelial cells and via TRPV4 in vascular smooth muscle cells [38,39]. These precedents, together with the well-established roles of TRP channels, in particular TRPA1, as regulators of cell signaling and their propensity to be activated by PM perturbations induced by various lipophilic factors, led us to hypothesize that these channels can be modulated by LNPs. Here, we tested this hypothesis by assessing whether TRPA1 function is altered by LNPs in an overexpression, as well as in a natural TRPA1 expression system.

## Results

### Implementation of an adapted Ca^2+^ -imaging setup

We used Fura2-based intracellular Ca^2+^ imaging to assess the acute effects of LNP preparations on TRPA1-expressing cells. To replicate more closely the *in vivo* scenario of slow particle diffusion through the tissue, we applied the test solutions via a custom-made slow injection system (see Methods and *Figure 1A*), followed by a period of no perfusion to allow the undisturbed interaction of the samples with the target cells. Then, the regular gravity-driven perfusion system was turned on to wash out the cells and to apply the channel agonist (100 μM AITC), which served to identify the TRPA1-expressing cells. We performed the experiments in a thermo-regulated chamber to ensure that the suspected particle-membrane-TRPA1 interactions took place at a physiological temperature of 35 ×C (see Methods and *Figure 1A*).

**Figure 1.**
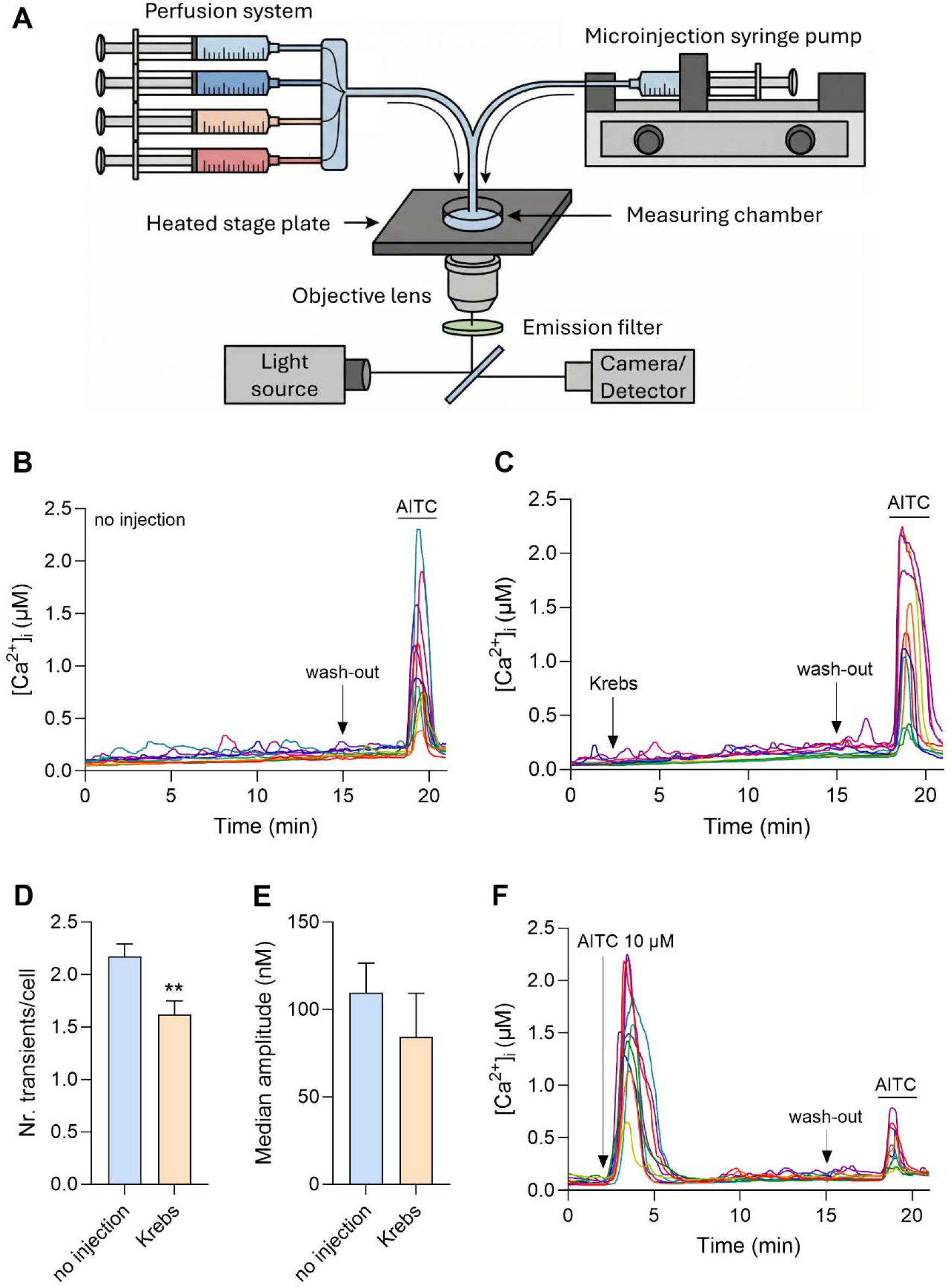
***A*** The experimental setup used for intracellular Ca^2+^-imaging experiments (schematic created with the website *deevid.ai*). **B** Representative traces of basal activity in CHO-mTRPA1 cells at pH 7.4 in the absence of any shear stress (no injection). **C** Representative traces of basal activity in CHO-mTRPA1 cells at pH 7.4 upon Krebs injection. **D**,**E** Quantification of basal activity Ca^2+^ responses in CHO-mTRPA1 cells at pH 7.4: (D) number of transients per cell and (E) median amplitude of the Ca^2+^ transients induced in responding cells. Asterisks correspond to the comparison between the two conditions: no injection vs. Krebs injection (P < 0.05). **F** Representative traces of Ca^2+^ responses elicited by the injection of 10 μM AITC in CHO-mTRPA1 cells at pH 7.4. AITC (100 μM) was applied at the end of the all experiments protocol, as positive control for TRPA1 expression.

CHO-mTRPA1 cells displayed small spontaneous Ca^2+^ transients in resting conditions (no injection or perfusion; *Figure 1B*). These transients were analyzed using a customized automatic algorithm that yielded the percentage of cells displaying at least one Ca^2+^ transient, the average number of Ca^2+^ transients per responding cell, and the median amplitude of Ca^2+^ transients in responding cells (see Methods). Similar Ca^2+^ transient activity was observed upon injection of control Krebs solution with the syringe pump (*Figure 1C-E*), indicating that the potential mechanical perturbations (shear stress) produced by this maneuver did not have significant effects on the intracellular Ca^2+^ dynamics. This basal activity was TRPA1-dependent, because it was fully blocked by the specific channel inhibior HC-030031 and did not occur in CHO-WT cells (*Figure 3D*). Injection of AITC (10 µM) induced responses in CHO-mTRPA1 cells showing a steep ascending slope followed by a slower decay due to channel desensitization, as illustrated by the weak responses to AITC at the end of the experiment (*Figure 1F*). These responses were similar to those produced by the classical continous perfusion (compare with the responses to AITC in panels B and C). This indicated that the modified setup was appropriate to assess TRPA1 responses.

### LNPs trigger cytosolic Ca^2+^ transients in CHO-mTRPA1 cells

We tested the effects of ionizable LNPs with a composition similar to that of LNPs currently licensed for therapeutics (55.5% DLin-MC3-DMA, 25% cholesterol, 13.5% DSPC and 6% DMG-PEG-2000) [40,41]. The LNP preparations and the control solutions (vehicle and Krebs) were characterized using the Nanoparticle Tracking Analysis (NTA, *Table 1*). This revealed that LNPs have a median diameter of 150 ± 7 nm.

**Table 1.** NTA characterization of LNP and control solutions used in the measurements.

|  | LNPs | vehicle | Krebs |
| --- | --- | --- | --- |
| Concentration (p/mL) | $3.69 \times 10^{12} \pm 1.62 \times 10^{12}$ | $8.05 \times 10^9 \pm 3.69 \times 10^9$ | $6.86 \times 10^6 \pm 1.84 \times 10^6$ |
| Median size (nm) | $149.97 \pm 6.77$ | $144.59 \pm 4.08$ | $191.05 \pm 16.80$ |
| Mean size (nm) | $171.97 \pm 8.65$ | $165.46 \pm 6.96$ | $214.83 \pm 13.75$ |

Application of a highly concentrated LNP suspension (10^11^ p/mL) at pH 7.4 increased the proportion of CHO-mTRPA1 cells showing Ca^2+^ transients and the number of transients per cell. However, these effects were similar to those induced by application of the vehicle (*Figure 2A, C-E* and *Figure S1*), suggesting that the LNPs did not induce any response on their own at this pH. Suspecting that the interaction of the LNPs with the PM is enhanced when their surface potential is positive, we tested the effects of LNPs (10^11^ p/mL) at pH 6.2 (below the pK_a_ value of 6.44 for the ionizable lipid). In this condition LNPs triggered large Ca^2+^ transients in CHO-mTRPA1 cells. These responses showed an unusual profile, compared to the typical shape of the responses to AITC. They appeared as one or multiple Ca^2+^ transients of different amplitudes throughout the application time (*Figure 2B*). This activation pattern may be a result of the stochastic interactions between the diffusing LNPs and the plasma membrane of the targeted cells.

**Figure 2.**
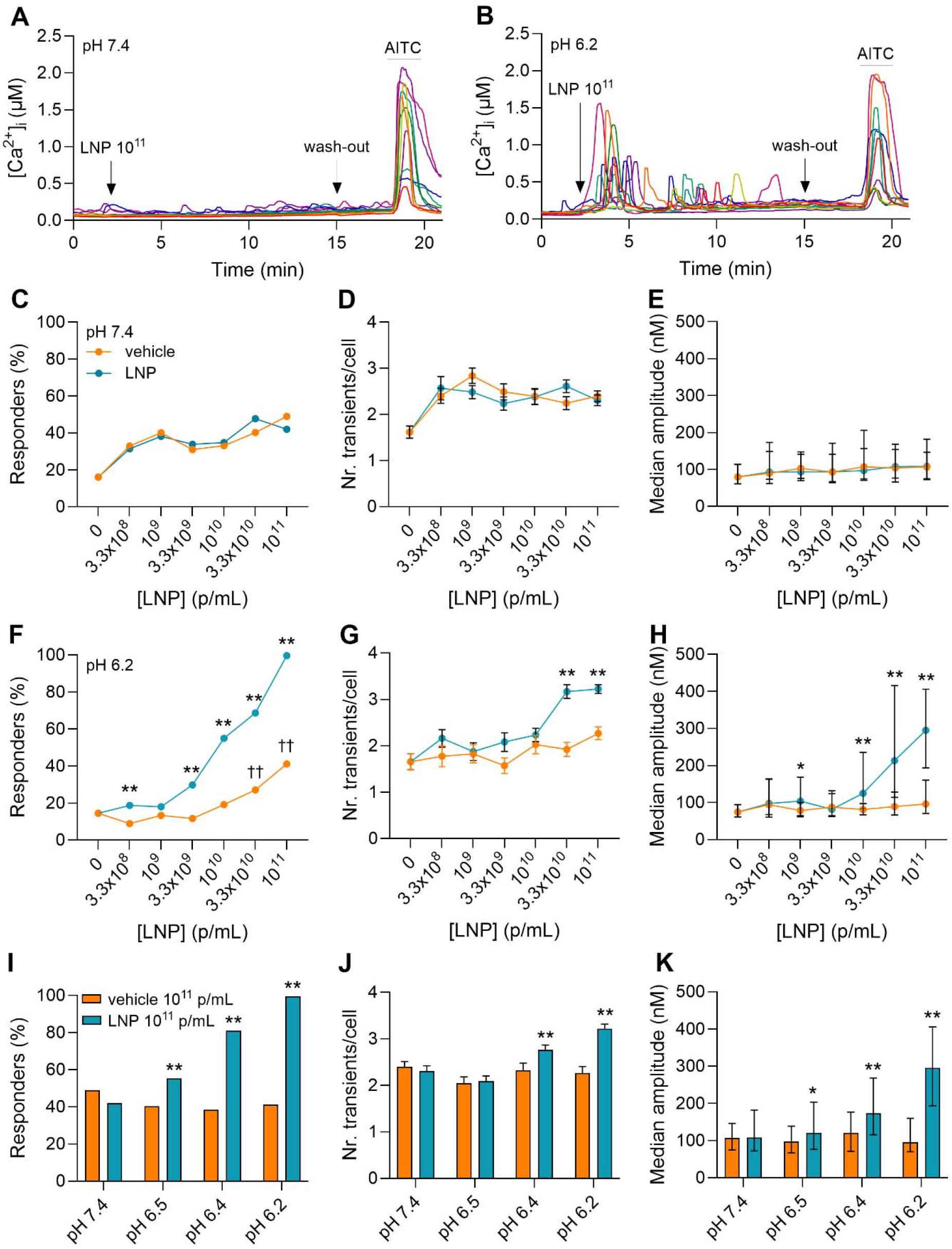
Quantification of Ca^2+^ responses induced in CHO-mTRPA1 cells by the application of LNPs and control solutions. **A** Representative traces of Ca^2+^ responses elicited by the injection of LNP 10^11^ p/mL in CHO-mTRPA1 cells at pH 7.4. **B** Representative traces of Ca^2+^ responses elicited by the injection of LNP 10^11^ p/mL in CHO-mTRPA1 cells at pH 6.2. **C-E** Dose response curves of the percentage of responding cells (C), number of Ca^2+^ transients (D) and median amplitude of Ca^2+^ transients (E) in the responding cells at pH 7.4. **F-H** Dose response curves of the percentage of responding cells (F), number of Ca^2+^ transients (G) and median amplitude of Ca^2+^ transients (H) in the responding cells at pH 6.2. **I-K** pH dependence of the percentage of responding cells (I), number of Ca^2+^ transients (J) and median amplitude of Ca^2+^ transients (K) induced by LNP 10^11^ p/mL. The asterisks (*) correspond to the comparison between each concentration of LNPs and the corresponding vehicle solution. The daggers (†) corresponds to the comparison between each concentration of vehicle solution and Krebs (LNP 0 p/mL).

Lowering the pH to 6.2 resulted in more pronounced responses that were dependent on the LNP dose, as reflected in all tested parameters (*Figure 2F-H*). Of note, the control (vehicle) solutions corresponding to the two highest doses (3.3 x 10^10^ and 10^11^ p/mL) recruit significantly more cells compared to the application of Krebs. However, this is not reflected in the number of transients per cell, nor in the median amplitude of the responses (*Figure 2G,H*). When assessing the effect of pH we found that the LNP-induced Ca^2+^ transients were most pronounced at pH values below the pK_a_ value of DLin-MC3-DMA (*Figure 2I-K*). Because the most noticeable effects are visible for the highest dose (10^11^ p/mL) and the lowest pH tested (6.2), where virtually all cells exhibit at least one Ca^2+^ transient, we used this condition throughout the rest of the study, unless stated otherwise.

### The LNP-induced Ca^2+^ responses are partly mediated by TRPA1

To test whether TRPA1 is involved in the LNP-induced generation of Ca^2+^ responses, we applied the LNP preparations in the presence of specific TRPA1 inhibitors (HC-030031 or A-967079) or the non-specific cation channel blocker ruthenium red (RR). All three blockers tested significantly reduced the LNP-induced Ca^2+^ responses in all three assessed parameters (*Figure 3A-C*). Representative Ca^2+^ responses induced by LNP preparations in the presence of HC-030031 or RR are presented in *Figure 3D,E*, respectively. Interestingly, the two specific TRPA1 blockers seem to have different effects on the time distribution of the Ca^2+^ transients throughout the application time. HC-030031 and A-967079 were more effective at blocking the responses in the second half of the application window, while allowing some of the early responses to occur. In contrast, RR is more effective at blocking the early Ca^2+^ responses (i.e. the first half of the application window) *(Figure S2)*. However, a full inhibition was not observed for any of these compounds. From the residual responses in the presence of the specific channel blockers it can be concluded that roughly 40-50% of the responses are TRPA1-independent. This suggests that TRPA1 is only partly involved in the generation of the Ca^2+^ transients induced by the LNP preparations, and other, yet unknown, pathways are also recruited by this stimulus. The non-specific blocker had a stronger inhibitory effects on the responses than HC-030031 and A-967079, indicating that at least part of the TRPA1-independent responses are mediated by Ca^2+^ influx from the extracellular medium.

**Figure 3.**
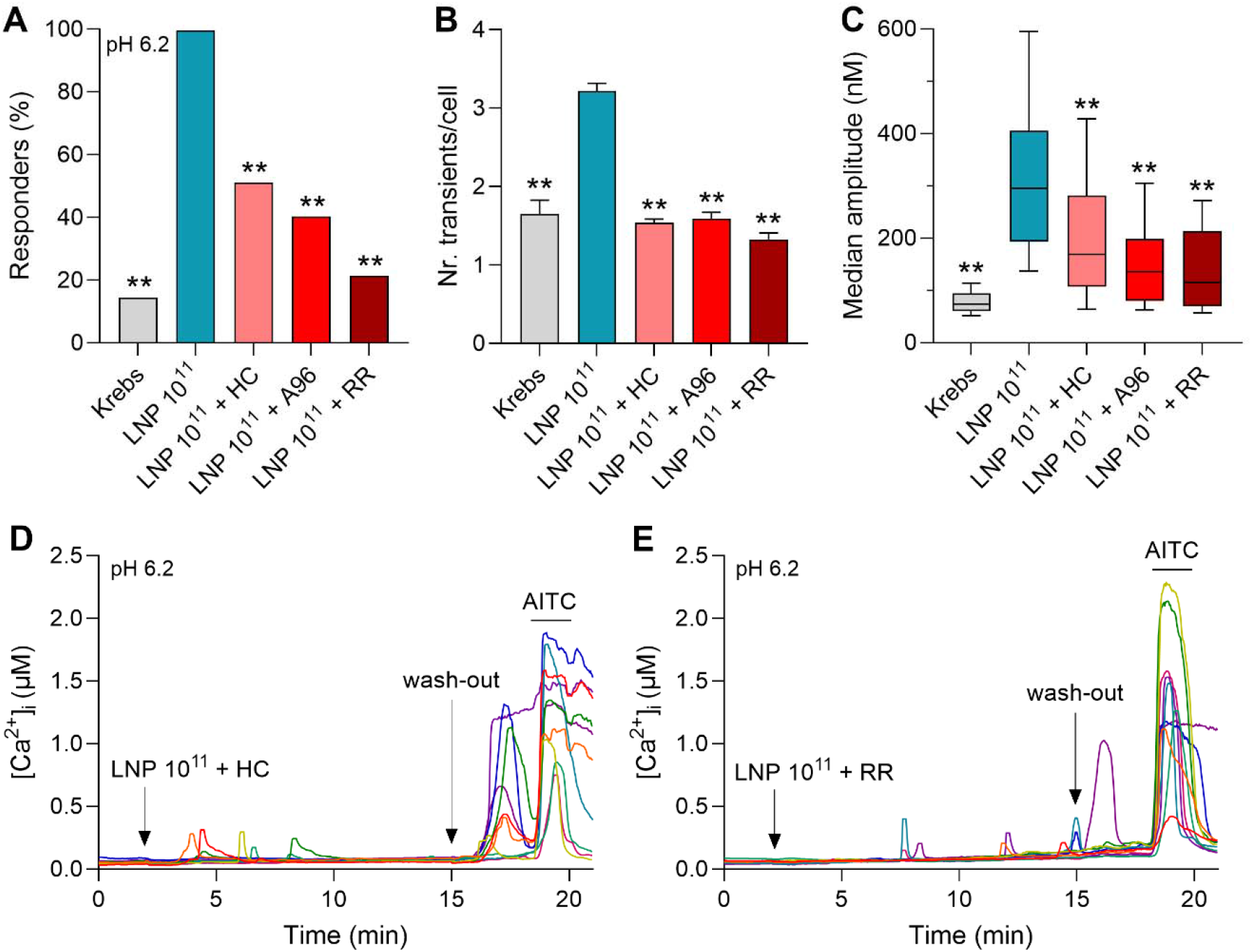
**A-C** Quantification of Ca^2+^ responses induced by LNP preparations in the presence of channel blockers in CHO-mTRPA1 cells at pH 6.2: (A) percentage of responding cells, (B) number of transients per cell and (C) median amplitude of the Ca^2+^ transients induced in responding cells. **D** Representative traces of Ca^2+^ responses in the presence of HC-030031. **E** Representative traces of Ca^2+^ responses in the presence of RR. The asterisks correspond to the comparison between each application and LNP 10^11^ p/mL.

### LNP preparations induce plasma membrane fluidization

Because lipid particles are roughly one order of magnitude larger than the TRPA1 channel, it is highly unlikely that these structures interact with binding pockets in the protein. Alternatively, LNPs can interact with the PM through surface adsorption, endocytosis and fusion, all of which involve PM mechanical perturbations. To test whether our samples alter lipid bilayer properties of CHO-mTRPA1 cells at pH 6.2, we used 1,6-diphenyl-1,3,5-hexatriene (DPH), a chemical probe serving to assess membrane fluidity by monitoring the fluorescence anisotropy [42]. The two highest LNP doses (3.3 x 10^10^ and 10^11^ p/mL) induced measurable decreases in fluorescence anisotropy in CHO-mTRPA1 cells, which are time-dependent and peak towards the end of the measuring window. This indicates that the LNP preparations fluidize the membrane. The lower concentrations (3.3 x 10^8^ – 10^10^ p/mL) do not induce changes that can be detected by this probe (*Figure 4A*). The LNP suspensions induced comparable membrane fluidization effects in CHO-WT cells (*Figure 4B*).

**Figure 4.**
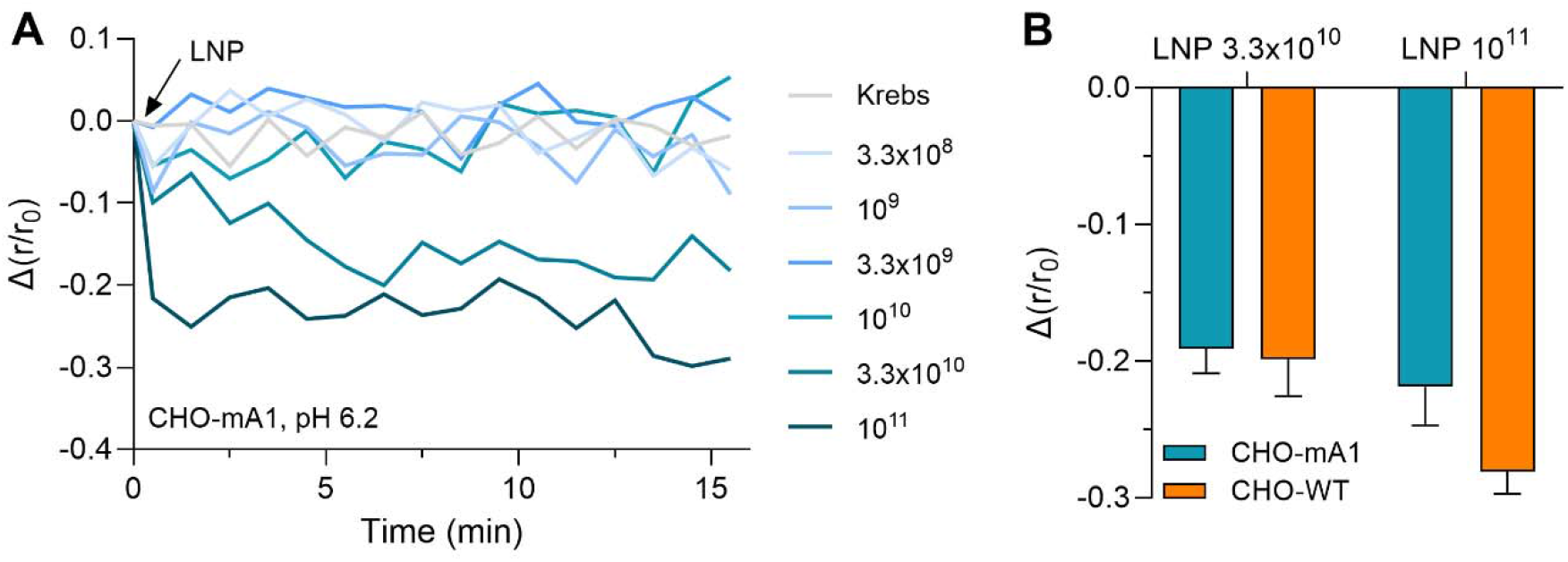
**A** Dynamics of relative fluorescence anisotropy measured in CHO-mTRPA1 cells exposed to increasing concentrations of LNP preparations at pH 6.2. **B** Quantification of the relative fluorescence anisotropy measured in CHO-mTRPA1 and CHO-WT cells at different time points.

### LNP preparations trigger cytosolic Ca^2+^ transients in CHO-WT cells

Since the channel blockers do not fully inhibit the responses induced by LNPs in CHO-mTRPA1 cells, we explored the possibility that these preparations might have other targets. For this, we performed the same type of Ca2+ measurements at pH 6.2 in CHO-WT cells, which lack endogenous TRPA1. Firstly, we found that the CHO-WT do not show basal activity upon the injection of Krebs (*Figure 5A*). Secondly, the application of concentrated LNPs resulted in transient Ca^2+^ responses (*Figure 5B*), but unlike CHO-mTRPA1 cells, the wild-type cells did not respond to the LNPs at low doses (3.3 x 10^8^ – 10^10^ p/mL) (*Figure 5C*). The Ca^2+^ transients induced in CHO-WT were less frequent than in CHO-mTRPA1 cells, with the highest concentration only inducing one-third of the responses elicited in CHO-mTRPA1 cells (*Figure 2I-K, Figure 5D-F)*. Moreover, no significant effects compared to control vehicle solutions were observed in terms of number of transients per cell and median amplitude of the transients in the case of responding CHO-WT cells, with the exception of the median amplitude corresponding to the highest concentration (*Figure S3*).

**Figure 5.**
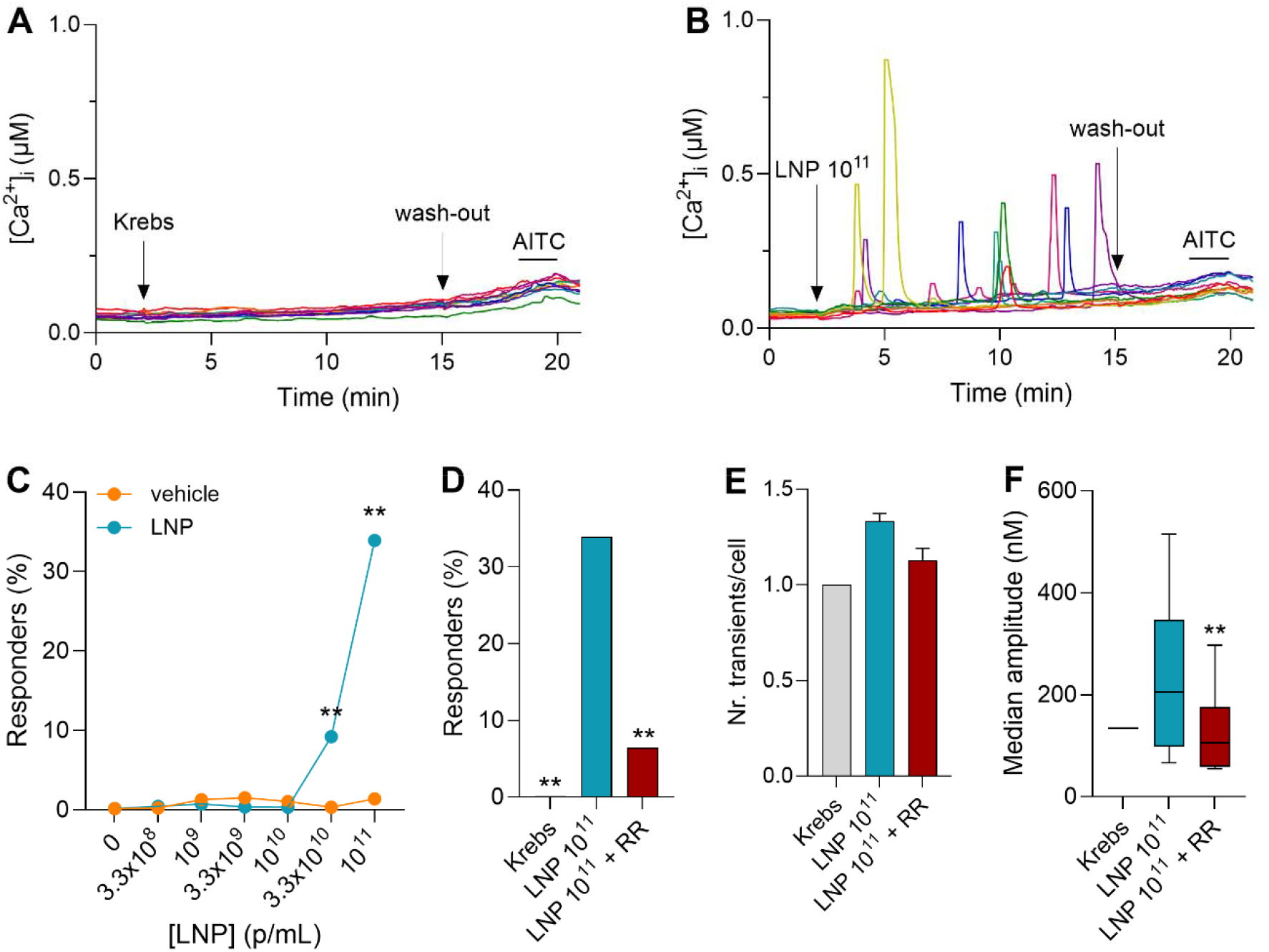
**A** Representative traces of basal activity in CHO-WT cells at pH 6.2. **B** Representative traces of Ca^2+^ transients induced by LNP 10^11^ p/mL in CHO-WT cells at pH 6.2. **C** Dose response of the percentage of cells responding to LNP 10^11^ p/mL in CHO-WT cells at pH 6.2. The asterisks correspond to the comparison between each concentration of LNP and the corresponding vehicle solution. **D-F** Quantification of Ca^2+^ responses induced by LNP preparations in the presence of the nonspecific Ca^2+^ channel blocker RR in CHO-WT cells at pH 6.2: (D) percentage of responding cells, (E) number of transients per cell and (F) median amplitude of the Ca^2+^ transients induced in responding cells. The asterisks correspond to the comparison between each application and LNP 10^11^ p/mL.

To test whether the increase in cytosolic [Ca^2+^] is a result of Ca^2+^ influx from the extracellular medium in CHO-WT cells, we used the non-specific Ca^2+^ channel blocker RR. This resulted in a 5-fold decrease in the number of responding cells and a 3-fold decrease in the median amplitude of the responses in activated cells, while the number of transients per cell remained unaltered (*Figure 5D-F*). The Ca^2+^ transients were strongly reduced by RR but not fully inhibited, indicating that LNPs may activate RR-insensitive Ca^2+^ influx systems and/or evoke Ca^2+^release from internal stores.

### LNPs release Ca^2+^ from the endoplasmic reticulum

When exposed to LNP preparations in the absence of extracellular Ca^2+^, 86% of CHO-mTRPA1 and 91% of CHO-WT cells exhibited cytosolic Ca^2+^ transients (*Figure S4*). This indicates that LNPs could evoke Ca2+ flux from intracellular Ca2+ stores, such as the endoplasmic reticulum (ER), which is the main Ca^2+^-storing organelle in cells. To test the possibility that LNP preparations trigger Ca^2+^ mobilization from the endoplasmic reticulum (ER), we employed two different approaches. Firstly, we directly measured Ca^2+^ levels in the ER of CHO-mTRPA1 and CHO-WT cells exposed to concentrated LNPs under low pH conditions, using the genetically encoded fluorescent ER Ca^2+^ indicator R-CEPIA1*er*. We observed a similar pattern of transient decreases in ER luminal [Ca^2+^] of multiple amplitudes, with one or multiple transients measured per cell (*Figure 6A,B*). In contrast, the application of Krebs did not elicit any Ca^2+^ transients in either cell type. Moreover, LNP preparations were able to mobilize ER Ca^2+^ to the same extent in the two cell types tested, as reflected by the lack of significant differences for the parameters tested (*Figure 6C-E*).

**Figure 6.**
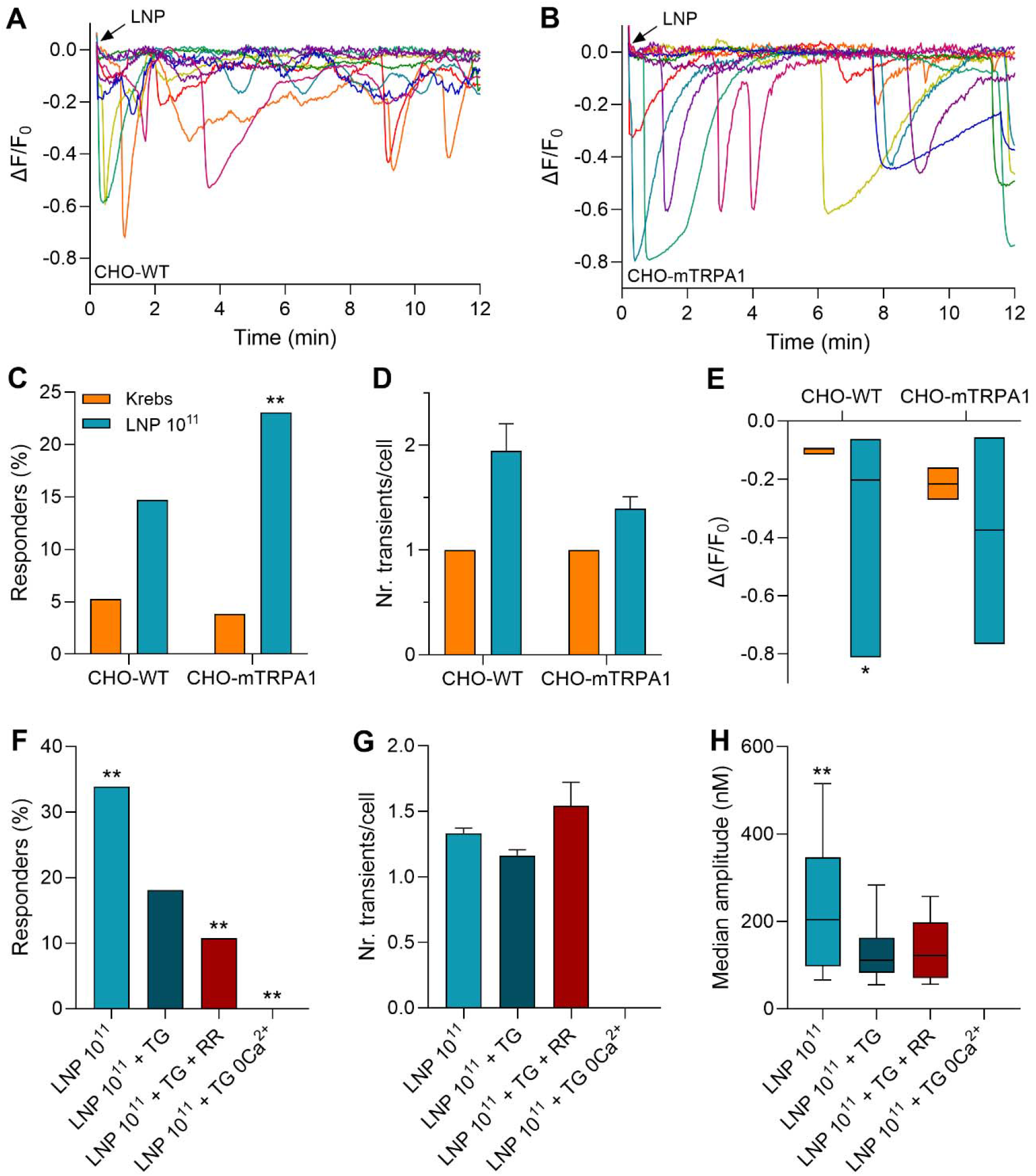
LNP preparations trigger Ca^2+^ mobilization from the ER. **A** Representative traces of ER luminal [Ca^2+^] in CHO-mTRPA1 cells at pH 6.2 upon injection of LNP 10^11^ p/mL. **B** Representative traces of ER luminal [Ca^2+^] in CHO-WT cells at pH 6.2 upon injection of LNP 10^11^ p/mL. **C-E** Quantification of the luminal Ca^2+^ release upon LNP application: (C) percentage of responding cells, (D) number of transients per responding cell, (E) median relative amplitude of the transients in responding cells. The asterisks correspond to the comparison between LNP 10^11^ p/mL and Krebs, for each cell line. **F-H** Quantification of the LNP-induced cytosolic Ca^2+^ transients upon ER emptying with thapsigargin: (F) percentage of responding cells, (G) number of transients per responding cell, (H) median relative amplitude of the transients in responding cells. The asterisks correspond to the comparison between each application and LNP 10^11^ p/mL + TG.

Secondly, to evaluate the contribution of ER Ca^2+^ mobilization on the cytosolic Ca^2+^ responses, we used thapsigargin (TG) to ensure full release of Ca^2+^ from the ER. TG is an irreversible, high-affinity SERCA pump inhibitor that prevents the reuptake of Ca^2+^ in the ER lumen. When CHO-WT cells were challenged with LNP preparations in the presence of TG, we observed a smaller population of responding cells, as well as smaller Ca^2+^ transients in the responding cells compared to the control conditions (application in the absence of TG). The average number of transients in the responding cells was not affected. The nonspecific Ca^2+^ channel blocker RR further reduced the population of responding cells, but left the other two parameters unaffected. This confirms the previous observation that LNPs are able to recruit a yet-to-be-identified RR-insensitive pathway that contributes to the Ca^2+^ transients in CHO-WT cells. Representative traces of Ca^2+^ responses induced by LNP upon ER depletion with TG can be observed in *Figure S5*. Since co-application of LNPs and TG in the absence of extracellular Ca^2+^ was not able to trigger any Ca^2+^ responses, we are sure that the residual, RR-insensitive Ca^2+^ transients are a result of Ca^2+^ influx via the PM, rather than the recruitment of other intracellular Ca^2+^ pools (*Figure 6F-H*). Since TRPA1 is a channel both activated and inactivated by Ca^2+^, we did not test the effects of ER emptying using TG in CHO-mTRPA1 cells due to the possible interference of a robust increase in cytosolic [Ca^2+^] with the TRPA1 channel activity.

### LNPs induce Ca^2+^ transients in DRG neurons

TRPA1 is present in a large subpopulation of sensory neurons, and sensory terminals are likely to encounter LNPs upon injection or inhalation of LNP-containing therapeutics. Thus, we tested the effects of our LNP preparations in a native TRPA1 expression system, namely primary cultures of mouse DRG neurons. Because in the previous set of measurements, LNPs triggered the clearest responses at low pH values, we performed all experiments in DRG neurons at the extracellular pH of 6.2. Here, we used a protocol similar to the one described above for CHO cells, with the only difference that after the wash-out period we applied agonists for the three main TRP channels involved in pain (PS for TRPM3, AITC for TRPA1 and capsaicin for TRPV1), in order to differentiate between different subpopulations of cells. We found that, in contrast to CHO cells, DRG neurons do not exhibit basal Ca^2+^ activity upon injection of Krebs solution (*Figure 7A*). In fact, the very low noise level allowed us to reduce the detection threshold for the Ca^2+^ transients to 25 nM instead of the 50 nM used for CHO cells. When we applied the LNP preparation (10^11^ p/mL), we observed the same pattern of activation as in CHO-mTRPA1 cells (*Figure 7B*). Of note, the LNP-induced activation in DRG neurons is much lower compared to CHO-mTRPA1 cells, with only 20% of neurons responding to the highest dose (vs. 100% in CHO-mTRPA1 cells). This was however a significant effect compared to the control vehicle solutions (*Figure 7C*). No noticeable differences were observed for the number of transients and median transient amplitude between LNP and vehicle preparations (*Figure 7D,E*). The lower responses could be attributed to the lower TRPA1-proteinlevels in these DRGs compared to the CHO-mTRPA1 overexpression system. To test the contribution of TRPA1 to these Ca^2+^ responses, we co-applied the LNP preparations with the two selective TRPA1 inhibitors HC-030031 or A-967079, or the non-selective cation channel blocker RR. All these blockers significantly reduced the number of neurons that responded to LNPs (*Figure 7F*). However, in line with the observations in CHO-mTRPA1 cells, none of them fully abolished the responses. No significant differences were observed for the other two parameters tested (*Figure 7G,H*).

**Figure 7.**
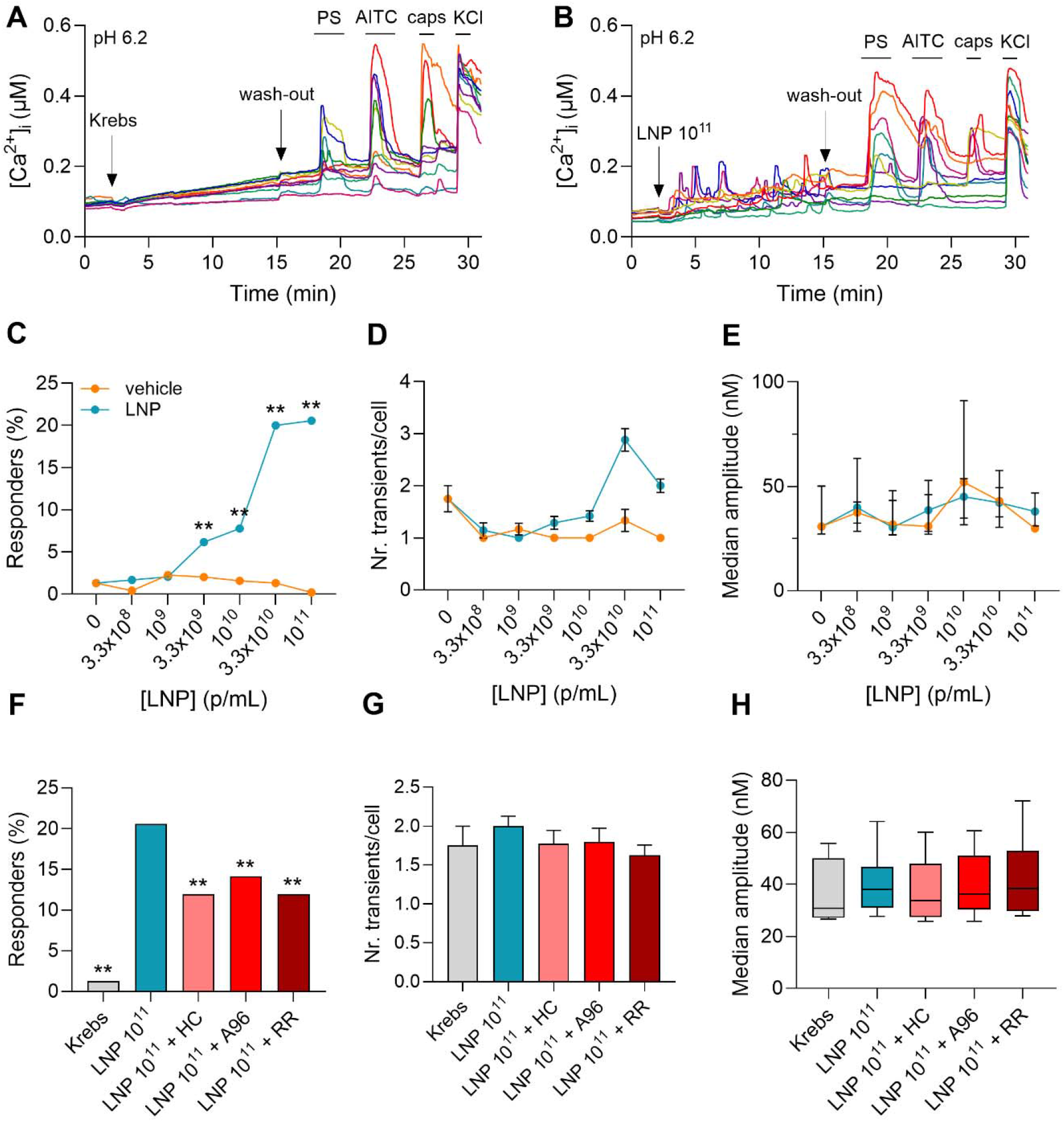
**A** Representative traces of basal activity in DRG neurons at pH 6.2. **B** Representative traces of Ca^2+^ transients induced by LNP 10^11^ p/mL in DRG neurons at pH 6.2. **C-E** Dose response curves of the percentage of (C) responding neurons, (D) number of Ca^2+^ transients and (E) median amplitude of Ca^2+^ transients in the responding neurons at pH 6.2. The asterisks correspond to the comparison between each concentration of LNP and the corresponding vehicle solution. **F-H** Quantification of Ca^2+^ responses induced by LNP preparations in the presence of TRPA1 channel blockers in DRG neurons at pH 6.2: (F) percentage of responding cells, (G) number of transients and (H) median amplitude of the Ca^2+^ transients induced in responding cells. The asterisks correspond to the comparison between each application and LNP 10^11^ p/mL.

DRG neurons are complex cell systems that express multiple TRP channels (TRPA1, TRPV1, TRPM3, TRPM8, etc.), as well as mechanosensitive ion channels (Piezo2), and multiple G protein-coupled receptors. It is thus possible that more than one actor is involved in mediating the responses to LNP preparations. We were interested in quantifying the LNP-induced Ca^2+^ transients in the subpopulation of DRG neurons that, out of the three TRP agonists tested, only responded to the application of AITC (i.e. they did not express measurable functional levels of TRPM3 and TRPV1). However, this population was very small, which did not allow us to perform reliable statistical tests (11% PS-/AITC+/caps-DRGs, out of which only 21% responded to the application of LNPs).

When we applied the LNP preparations in the absence of extracellular Ca^2+^, only a very small population of DRG neurons (3.5%) showed Ca^2+^ responses. This is in strong contrast with the much larger percentages of responding CHO cells (86% for CHO-mTRPA1 and 91% for CHO-WT). This is not surprising, since CHO cells and sensory neurons engage different Ca^2+^ mobilization pathways that rely on a different set of receptors. Because of the virtually absent Ca^2+^ mobilization in DRG neurons, we decided to not further explore the contribution of this pathway to the LNP-induced Ca^2+^ transients. Taken together, our results show that LNPs induce Ca^2+^ transients in DRG neurons. As in CHO-mTRPA1 cells, these responses are partly mediated by TRPA1, and that other, yet unidentified mechanisms are also involved in the generation of responses to LNPs.

## Discussion

Here, we tested the ability of ionizable LNP preparations to modulate the activity of TRPA1. For this, we used an LNP formulation similar to the currently approved LNP-based therapeutics: 55.5% DLin-MC3-DMA, 25% cholesterol, 13.5% DSPC, 6% DMG-PEG-2000 [40]. We used the lipid film hydration method to create the LNPs and we characterized our samples using a nano-tracking assay (NTA). The LNPs in our samples have a median size of 150 ± 7 nm. The LNPs used in COVID-19 vaccines and Onpattro are smaller (80-100 nm). Commercial LNPs are typically prepared using microfluidic systems, which allow for obtaining smaller particles compared to the manual extrusion through polycarbonate filter method [43,44]. We could not find any information about the concentration (particles/mL) of the licensed LNP-based therapeutics. However, the total lipid per dose is reported. For a comparison between the approved LNP-based therapeutics and the LNP preparations used in this study, please see *Table 2*. For the same amount of total lipid, the particle size is inversely proportional to the total number of particles. Considering that in the known LNP therapeutics the total lipid amount per volume is larger than in our preparations, and the size of the particles is smaller than ours, the other LNP-based therapeutics are expected to contain more particles per volume than the samples we tested.

**Table 2.**
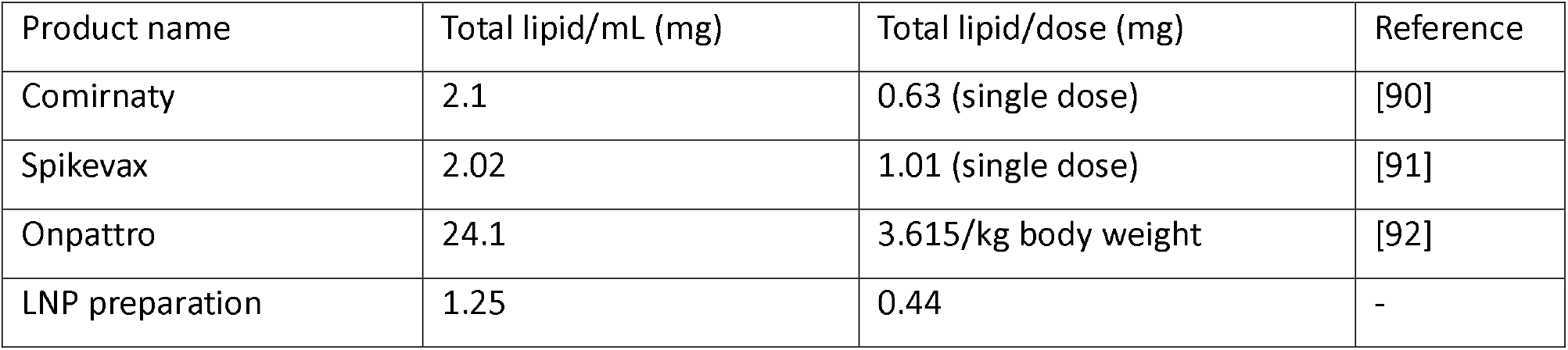
Comparison between the licensed LNP-based therapeutics and the LNP preparations used in this study.

To determine the effects of our LNP preparations on the TRPA1 channel, we adapted the Ca^2+^-imaging setup to allow the measurement in closer-to-physiological conditions – namely, slow particle perfusion, followed by resting, no-flow conditions, and measurement at physiological temperatures. Upon Krebs injection, CHO-mTRPA1 cells display a TRPA1-dependent basal Ca^2+^ activity, characterized by shallow, undulatory baselines. It is known that TRPA1 can be activated and inactivated by the pore-permeating Ca^2+^ upon stochastic channel opening [45]. The short-lived Ca^2+^ microdomains formed locally around the channel pore opening are amplified in a feedback loop that is terminated as a result of quick channel inactivation at high Ca^2+^ concentrations (low μM) [46–48]. The interaction between TRPA1 and Ca^2+^ is complex, as TRPA1 can directly bind calmodulin (CaM) under physiological conditions [49,50]. When Ca^2+^ is bound to CaM, CaM binding to TRPA1 promotes channel activation in low Ca^2+^ levels, while at high Ca^2+^ conditions, CaM binding to TRPA1 promotes channel inactivation [51–53]. The dynamic interaction between Ca^2+^, CaM and TRPA1 describe a self-regulating system in which a rise in intracellular Ca^2+^ leads to TRPA1 activation, followed by a quick inactivation once high levels have been reached. This interplay is achieved locally, in TRPA1-neighboring microdomains.

We used the customized experimental setup to measure the cytosolic Ca^2+^ responses in CHO-mTRPA1 cells upon controlled LNP injection. We determined that ionizable LNP preparations induced a dose- and pH-dependent response, in line with the protonation of the ionizable lipid at pH values below its pK_a_ value. Ionizable LNPs in the licensed therapeutics are designed to be electrically neutral until reaching the acidic environment of the endosome, where the protonation of the ionizable lipid causes the disruption of the endosomal membrane and endosomal escape. However, there are situations in which ionizable LNPs can encounter acidic tissues, such as the tumor microenvironment or the nasal mucosa. These two examples are relevant in the perspective of current therapeutic efforts to develop cancer and nasal vaccines. Reported pH values in these tissues can reach levels below the pK_a_ value of the ionizable lipid used in this study. In fact, the pK_a_ value represents the pH at which 50% of the ionizable headgroups have been protonated [54]. This effect reflects in the cell activation observed at intermediate pH values (*Figure 2I-K*). Besides their application as drug delivery systems, LNPs also have intrinsic adjuvant activity, that is superior to those of classic adjuvants such as aluminum salts and liposomes [55–57]. The adjuvant effect on the innate immune activation, characterized by cytokine production causes an inflammatory response that can lead to an acidic microenvironment [58], which may affect the protonation state of LNPs at the injection site. LNPs are trafficked from the injection site to the lymph nodes, where the pH can reach values of 6.2 – 6.8, low enough to protonate the LNPs [59]. Considering all the acidic environments mentioned above, the pH at which the measurements in this study was performed is of physiological relevance.

Immediately after injection, LNP preparations induced an atypical pattern of activation, characterized by one or multiple Ca^2+^ transients of different amplitudes, variably distributed throughout the time window of the measurements (12 min). This is in contrast with the classic TRP channel agonist response, characterized by a relatively smooth ascending slope, followed by a slower descent upon channel desensitization (*Figure 1F*). It is possible that this pattern of activation is a result of stochastic LNP-target cell interactions, as individual (or groups of) LNPs cause membrane perturbations during internalization. Under physiological conditions, LNP uptake by target cells is mainly mediated by endocytosis, and fusion events are rare. A study using LNPs of a similar composition, albeit of smaller size, shows that at neutral pH, LNP uptake mainly takes place by clathrin-mediated endocytosis and macropinocytosis and that internalization peaks around 6 h after exposure [60,61]. At 12 min, which is the time window used in our experiments, less than 0.5% of LNPs are internalized [62,63]. This is in line with our results showing no Ca^2+^ activity above the basal level in cells treated with LNPs at pH 7.4. On the other hand, a lower pH ensures ionizable lipid protonation and a much faster LNP-PM interaction based on electrostatic attraction between the cationic LNPs and the anionic membrane lipids. This leads to PM destabilization via the formation of a hexagonal H_II_ phase [64–66].

An important part of the LNP composition are PEG-lipids. After *in vivo* administration, the gradual dissociation of PEG-lipids from the LNPs ensures longer circulation times and targeting of specific tissues in the body [67]. In the case of approved LNP therapeutics, the short C14 PEG-lipids ensure a quick (within a few minutes) PEG dissociation and replacement by a biomolecular corona composed of electrolytes, proteins and lipids. *In vivo*, the biomolecular corona plays crucial roles in LNP fate by enabling recognition by specialized receptors and by triggering endocytosis. The formation of the protein corona is a major challenge in gene therapy and vaccine development, as it confers LNPs a new biological identity and alters their physicochemical properties [68–70]. In this study, the LNPs used have not been exposed to biological fluids before being applied to the cells. It is worth pointing out that the preservation of PEGylation could result in our LNPs to have physicochemical properties different from those of particles decorated with a biomolecular corona (e.g. altered surface charge, apparent size, shielding effects or promoting specific interactions with the target cells). Since our focus is on acute (<15 min) interactions with the target cells, the effects of PEGylation and of the corona are smaller in contrast with more prolonged (>1 h) effects, when LNPs are believed to have fully exchanged the PEG layer to a biomolecular corona. However, these effects should be bared in mind for the interpretation of the results of this study.

Within one minute after the injection, LNPs induce Ca^2+^ transients in target cells. The Ca^2+^ responses were reduced in the presence of multiple channel blockers, however not fully abolished, indicating only a partial involvement of TRPA1 in the generation of the Ca^2+^ transients. The differential effect of the specific and non-specific TRPA1 channel blockers on the time distribution of Ca^2+^ transients throughout the application time (i.e. the propensity to block the early vs. late responses) may be an indicative of different mechanisms involved in the occurrence of LNP-induced Ca^2+^ responses. The specific channel blockers HC-030031 and A-967079 are more effective in abolishing the late Ca^2+^ responses, but are also able to reduce the amplitude of the early responses, suggesting an involvement of TRPA1 throughout the entire time window of our experiments. On the other hand, the non-specific blocker of cation-permeable pores RR is more effective is abolishing the early responses (*Figure S2*). This is an indication that additional TRPA1-independent mechanisms are being engaged in the very acute phase of the exposure, such as other ion channels or LNP-induced Ca^2+^ pores in the plasma membrane. We confirmed the existence of a TRPA1-independent mechanism in CHO-WT cells, which lack endogenous TRPA1. Application of the same LNP preparations triggered Ca^2+^ transients in the wild-type cells, albeit to a significantly lesser extent. The CHO-WT responses were partially blocked by the non-specific Ca^2+^ channel blocker RR. Of note, RR is not able to cross the PM due to the large size, and the large positive electrical charge. This indicates that at least part of the Ca^2+^ responses arise as influx from the extracellular medium, either via other Ca^2+^-permeable channels, or via de novo-induced pores in the plasma membrane. *In silico* simulations with DLin-MC3-DMA in a protonated state (similar to our experiments) show that protonated LNPs can induce pores in the biological membranes they interact with. LNPs have aqueous pockets inside the lipid core that can be released in the cytosol upon membrane fusion [71]. In our case, the saline solution in which LNPs have been formed and stored contains 1.5 mM Ca^2+^. It is possible that the simultaneous release of several of these pockets is partly responsible for the Ca^2+^ transients observed. Moreover, the rise in local cytoplasmic Ca^2+^ concentration around TRPA1 could activate the channel, allowing for additional Ca^2+^ influx from the extracellular medium (since concentrations of 1 μM are enough to trigger TRPA1 activation). However, this effect is most likely too small to be the main mechanism by which LNP induce Ca^2+^ transients in target cells. It cannot be excluded that LNP form multiple kinds of cationic pores at the PM level.

We next tried to elucidate the mechanism by which LNPs induce the unusual activation pattern, characterized by Ca^2+^ transients, in CHO-mTRPA1 cells. Previous studies from our group have established that TRPA1 is sensitive to the properties of the local lipid environment, and plasma membrane perturbations can induce TRPA1 activation. In order to test this hypothesis, we used the fluorescent membrane probe DPH, which reports on the fluidity of the membrane. We noticed that only the two highest LNP concentrations used caused a decrease in DPH fluorescence anisotropy, indicating comparable membrane fluidization in both CHO-mTRPA1 and CHO-WT cells. This is in line with the dose-dependent Ca^2+^ responses in CHO-WT, where we show that only the two highest concentrations of LNP are able to induce measurable Ca^2+^ transients. It is possible that at such high concentrations, the membrane integrity of the cells is compromised by the LNP-PM interactions, allowing for ion exchange between the extracellular and intracellular environments. Additionally, the altered PM mechanical properties may also contribute to the TRPA1-mediated Ca^2+^ responses in CHO-mTRPA1 cells. Of note, previous studies on PM-inserting molecules such as alkylphenols and bacterial LPS point in the direction of an acute decrease in lipid bilayer fluidity upon treatment application [10,11]. This is in contrast with the current results, and could be a consequence of the fact that, unlike the small lipophilic molecules that insert in the membrane, LNPs do not interact preferentially with the hydrophobic lipid tails, but with the whole plasma membrane.

When applied in the absence of extracellular Ca^2+^, LNP preparations remained capable of transiently elevating cytosolic [Ca2+], indicating LNPs can evoke Ca^2+^ mobilization from internal stores (in particular the ER). This has been previously shown in another study, in which multiple cationic liposomes induced PLC-dependent Ca^2+^ mobilization, but no mechanistic explanation was provided [72]. However, the Ca^2+^ measurements in that study do not have single-cell resolution; thus it remains unknown whether cationic liposomes induce a similar pattern of transient Ca^2+^ responses. The mechanism underlying Ca^2+^ mobilization by LNPs is intriguing. Although our study does not provide sufficient information to draw conclusions about the pathways involved in this process, we envisage two possible explanations: LNPs induce PLC-dependent Ca^2+^ release (either GPCR-dependent or -independent), or LNPs induce perturbations in ER membrane. For the first scenario, it is possible that yet-to-be-identified G-protein coupled receptors are activated by LNPs via an unknown mechanism, leading to G_q_ protein activation, and subsequent PLC-dependent activation of IP_3_Rs in the ER. Alternatively, the possibility of GPCR-independent PLC activation cannot be excluded. Although not yet described for PLC, mechanical activation of other enzymes docked at the PM has been described. For example, diacylglycerol kinase (DGK) can be activated by high membrane curvature [73], while mechanical disruption of lipid rafts leads to phospholipase D (PLD) activation [74]. Alternatively, LNPs may alter the properties of ER membranes similarly to the mechanical alterations induced in the PM. It is well known that the ER is a widespread organelle, with multiple PM contact sites, where the ER membrane comes in very close proximity to the PM [75]. Following lipid exchange between the ER and cellular membranes, which has already been described [76], the H_II_ hexagonal phase induced by LNPs could also be induced in the ER membrane, compromising its integrity. In order to better understand the ability of LNP-induced Ca^2+^ mobilization from internal stores and, in particular the ER, further studies are needed.

Due to its activation by intracellular Ca^2+^, TRPA1 can support a positive feedback loop that amplifies small initial Ca^2+^ responses. Here, we observed that in CHO-mTRPA1 cells specific and non-specific channel blockers reduce the cytoplasmic LNP-induced Ca^2+^ responses, but do not block them fully. The quantification of Ca^2+^ transients in the two cell types shows that LNP recruit more cells and induce more frequent responses of higher amplitude in CHO-mTRPA1 than in CHO-WT cells. This observation is an indication that, in CHO-mTRPA1 cells, the transient Ca^2+^ release events from the ER are amplified by the channel, leading to more sustained responses compared to CHO-WT. This hypothesis is also supported by the fact that, in DRG neurons, where LNP trigger minimal Ca^2+^ mobilization from internal stores (3.34% of all neurons) (*Figure S4C,F)*, the TRPA1 blockers also have a much smaller, non-significant effect on the Ca^2+^ responses (*Figure 7F-H*), independent of the functional expression of TRPA1.

Lastly, we investigated the effects in LNPs in a native TRPA1 expression system: primary cultures of sensory DRG neurons. This is relevant in the context of LNP delivery, since in many cases, the administration of these therapies is linked to a pain-related context (e.g. cancer-related pain, vaccine-induced pain, etc.). Notably, LNPs induced a similar pattern of transient Ca^2+^ responses as in CHO-mTRPA1 cells, albeit at a much smaller extent. The Ca^2+^ mobilization from internal stores was almost absent, probably due to different molecular machineries and Ca^2+^ release dynamics compared to CHO cells (i.e. IP_3_R- or RyR-dominant) [50,77,78]. When we assessed the involvement of TRPA1 in the formation of Ca^2+^ transients using specific and non-specific TRPA1 channel blockers, we observed a significant reduction in the percentage of responding cells, but no change in the number of transients per cell or the median amplitude of the transients. Of note, CHO-mTRPA1 is an overexpression system and TRPA1 levels are much higher compared to the native expression in DRG neurons. One limitation of this study is testing the effects of LNPs on DRG cell bodies, opposed to nerve terminals, as the expression of plasma membrane receptors and, in particular, TRPA1 could differ considerably between these two regions of the neuron. TRPA1 is a sensor of noxious environmental stimuli and, from a functional perspective, the expression at the level of the nerve terminals is more relevant for pain signaling than at the level of the cell body located deep in the spinal cord. However, to the best of our knowledge, no studies have yet compared the expression of TRPA1 in different parts of the DRG neurons, and investigating the effects of LNPs on nerve terminals is particularly challenging given the constrained experimental conditions required here. Another important aspect when considering the interaction of LNPs with nerve terminals, besides the possible differential expression of ion channels, is the increased plasma membrane curvature. It is known that high membrane curvature promotes lipid membrane fusion [79] and that highly curved membranes promote endocytosis in a clathrin-independent manner [80]. This points in the direction of an increased LNP uptake at the level of sensory nerve terminals, compared to cell bodies. Moreover, as a future perspective, it would be relevant, for the context of pain, to investigate the ability of LNPs to induce action potentials in sensory neurons, as well as the role of LNPs in the modulation of the firing pattern induced by different stimuli.

To conclude, here we show that ionizable LNPs induce irregular Ca^2+^ transients in target model and native cells (CHO and DRG neurons), both dependently and independently of TRPA1. However, the effects of LNPs in CHO-mTRPA1 cells are significantly more pronounced than in CHO-WT cells. We envisage that LNPs induce a multifactorial effect in CHO-mTRPA1 cells. Firstly, LNPs induce Ca^2+^ mobilization from internal stores, in particular the ER. Secondly, LNPs trigger Ca^2+^ influx across the plasma membrane. Part of the Ca^2+^ influx is reduced in the presence of TRPA1 channel blockers, indicating a partial contribution of TRPA1 to the observed Ca^2+^ transients. The TRPA1-mediated Ca^2+^ influx can be partly produced downstream of Ca^2+^ mobilization from the ER and acts in amplifying the Ca^2+^ release from internal stores. Other direct effects, such as TRPA1 activation induced by plasma membrane fluidization or direct activation by trace free lipids in the LNP preparations could also contribute to the TRPA1-dependent Ca^2+^ responses. More work is required to identify the molecular mechanisms underlying the remaining TRPA1-independent Ca^2+^ influx induced by LNPs. These particles may engage other players at PM, such as other ion channels and Gq-protein coupled receptors, or could directly induce Ca^2+^-permeable pores in the PM. Nonetheless, here we have identified a new class of TRPA1 activity modulators, lipid nanoparticles, that induce a non-canonical pattern of activation. These results serve as a starting point in a better understanding the interplay between TRP channels and various lipid particles involved in physiological and pathophysiological processes in the body.

## Methods

### Cell culture

Chinese hamster ovary cells (RRID:CVCL_0213) (denoted CHO-WT) from the American Type Culture Collection were grown in DMEM containing 10% fetal bovine serum, 2% glutamax (Gibco/Invitrogen), 1% non-essential amino acids (Invitrogen) and 200 µg/mL penicillin/streptomycin at 37 ×C in a humidity-controlled incubator with 5% CO_2_. As mTRPA1 expression system we used CHO-K1 cells (RRID:CVCL_0214) stably transfected with mTRPA1 (denoted CHO-mTRPA1). The cells have been seeded on poly-L-lysine-coated 13 mm coverslips 2h before the measurements and cultured in 24-well plates.

### Microinjection system and temperature control

We implemented several modifications in the setup to replicate more closely the *in vivo* scenario of slow particle diffusion through the tissue. We replaced the classic, constant perfusion of the cells with a gentle injection of test solutions,

For this, we used a microinjection syringe pump (Fusion 200 precision syringe pump, Chemyx, Stafford TX, USA) to perfuse 350 μL LNP suspension over 1 min on top of an equal volume of Krebs solution (already in the measuring chamber). In a typical setup, this is achieved by perfusing the test solutions via a Peltier heating element. Since this arrangement does not allow attaching a Peltier element to the syringe pump, a heated plate holder mounted on the microscope table had to be implemented to allow for a constant heating of the cells at a set temperature.

### Ratiometric intracellular Ca^2+^ imaging

For intracellular Ca^2+^ imaging experiments, the cells were incubated with 6 µM Fura-2 AM (Biotium, Hayward, CA, USA) for 60 min at 37 °C in a 5% CO_2_, humidity-controlled incubator. CHO-WT were co-incubated with Pluronic F-127 20% solution in DMSO (1:40 dilution, Biotium) to facilitate cell loading. Fluorescence was measured with alternating excitation at 340 and 380 nm using a monochromator-based imaging system consisting of an MT-10 illumination system (Tokyo, Japan) and Cell-M software from Olympus. All experiments were performed at 35 °C using a standard Krebs solution. Fluorescence intensities were corrected for background signal and converted to intracellular Ca^2+^ concentrations.

Only the cells that responded to the positive control (AITC) at the end of the experiment were considered TRPA1-expressing cells and included in the analysis.

### Measurements of ER Ca^2+^

ER Ca^2+^ levels were measured using a genetically encoded Ca^2+^ indicator R-CEPIA1*er*, previously developed by the team of Dr. M. Iino [81]. The constructs were introduced in the cells using the Lipofectamine 3000 transfection reagent (Sigma-Aldrich), according to the manufacturer’s indications. For the measurements performed in the absence of ER Ca^2+^ mobilization, the cells have been incubated with thapsigargin for 5 min before the application of LNP preparations. This resulted in the immediate release of luminal Ca^2+^, followed by a return of cytosolic Ca^2+^ to the baseline.

### Preparation of ionizable LNPs

The ionizable LNPs were prepared using the thin lipid film hydration technique, followed by water bath sonication and polycarbonate membrane extrusion (Mini Extruder, Avanti), as previously described [82]. In short, 25 mg/mL lipid stocks were prepared by dissolving lipids in deuterated chloroform (Sigma-Aldrich) and were mixed in a glass tube, then dried under constant nitrogen flow until a film formed on the surface of the tube. The lipid film was further dried overnight in a vacuum desiccator. The next day, the lipid film was rehydrated using hot (95 ×C) standard Krebs solution. The LNP suspension was sonicated to decrease the vesicle size. The solution was passed via polycarbonate filter with pore sizes of 400 nm, followed by 100 nm. The LNP preparation was stored at 4 ×C. The composition of the LNPs is similar to that of the currently licensed LNP-based therapeutics: 55.5% DLin-MC3-DMA (MedChem Express), 25% cholesterol (Sigma Aldrich), 13.5% DSPC (Sigma Aldrich), 6% DMG-PEG-2000 (Sigma-Aldrich). For the vehicle control solutions, deuterated chloroform without lipids was subjected to the same protocol steps. The preparations have been characterized for their size and concentration by a nanoparticle tracking assay (NTA). For this, the concentrated samples were pre-diluted and analyzed using a ZetaView analyzer. Samples were measured at 11 different positions in the chamber, with stage temperature control set at 22 °C. After the NTA measurement, the stock solution was diluted to a storage concentration of 10^12^ particles/mL. The different concentrations used in the measurements were obtained via serial dilution from the main stock.

### Identification of [Ca^2+^]_i_ peaks and quantification of [Ca^2+^]_i_ transients

In order to quantify the Ca^2+^ responses to LNP preparations, we implemented an automated analysis algorithm that: 1) identifies the baselines of the traces and corrects for the drift that arises as a result of fluorophore bleaching, 2) identifies the peaks in the traces using a dual threshold (a dynamic threshold based on the noise level of the recording and a static value of 50 nM for CHO cells and 25 nM for DRG neurons, 3) for each peak identified, it records the time of occurrence and the amplitude above the trace baseline level, and 4) for each cell, it records the number of peaks that meet the threshold criteria. Firstly, a second-order polynomial function is fitted at the bottom of the recorded traces in order to “flatten” the recordings and correct for any unwanted drift. A description of the algorithm used in our analysis (the modified polynomial algorithm) can be found here [83]. Afterwards, the corrected traces are further used to detect and quantify the Ca^2+^ transients. In short, the peak detection algorithm used for the identification of Ca^2+^ transients finds local maxima by comparing each point-value to the values of the neighboring data points within a window of desired width. If the prominence of the peak relative to its surrounding baseline exceeds a given dynamic threshold (based on the noise level of the recording), the peak is considered for further analysis. If the time distance between two consecutive peaks is smaller than a given value, the second peak is not taken into consideration. A description of the peak detection algorithm used can be found here [84,85]. The adapted Pyhton script used for analysis is freely available on GitHub [86].

### Fluorescent measurements using DPH

A 1 mM stock solution of 1,6-diphenyl-1,3,5-hexatriene (DPH) (Sigma-Aldrich) was prepared in dimethyl sulfoxide (DMSO) and a working solution of 10 µM was prepared in Krebs. Next, 1.5 µM DPH was prepared in standard Krebs solution to stain CHO-mTRPA1 cells (1 × 10^6^ cells/mL) for 30 min at 37 ×C. After incubation, the cells were washed with Krebs, and suspensions of 100 μL were aliquoted into flat-bottom 96-well microtiter plates (Greiner Bio-One, Vilvoorde, Belgium). DPH anisotropy (r) was monitored with an excitation wavelength of 365 nm and an emission wavelength of 430 nm using a Flexstation III Benchtop Multi-Mode Microplate Reader and the SoftMax Pro Microplate Data Acquisition & Analysis Software (Molecular Devices). A linearly-polarized excitation beam was generated by a vertical polarizer that excites DPH with transition moments aligned parallel to the incident polarization vector. The resulting fluorescence intensities of emitted light in both parallel (I_VV_) and perpendicular (I_VH_) directions to that of the excitation beam were recorded and the fluorescence anisotropy calculated by: r = (I_VV_ − G·I_VH_)/(I_VV_ + 2G·I_VH_), where G = I_HV_/I_HH_ [87].

### Isolation and culture of dorsal root ganglion (DRG) neurons

Primary cultures of DRG neurons were obtained from C67BL/6 wild-type mice (RRID:IMSR_JAX:000664) (12 weeks old, mixed sexes) following a protocol similar to the one described here [88]. In short, animals were euthanized in a CO_2_ chamber right before the dissection. Incisions were made under both scapulae and the neck to reach the spine. The spine was extracted by detaching it from the underlying tissue. After extraction, the spine was cleaned from any remaining muscle, tissue, and excess bone fragments. The cleaned spine was bisected, and the spinal cord was carefully removed with forceps to avoid disturbing the dural sheath membrane covering the DRGs. Bisected halves were stored on ice in a Petri dish containing PBS with penicillin and streptomycin (3%). The dural sheath covering the DRGs was removed, and the DRGs were carefully extracted using the forceps. Extracted DRGs were temporarily kept on ice and cleaned by removing the attached axons. Finally, the cleaned DRGs were stored in a 1.5 ml Eppendorf tube with basal medium (BM) (90% neurobasal medium (NBM), 10% fetal calf serum). The dissociation of DRG neurons involved two steps: chemical dissociation using 2.5 mg/mL dispase and 2 mg/mL collagenase (incubation at 37°C for one hour), followed by mechanical dissociation (via 22G and 26G syringe needles). This cell suspension was carefully layered on top of 3 ml of 16% bovine serum albumin (BSA) solution and centrifuged for 6 min at 500 x g to filter out the debris. The supernatant was removed, and the pellet was resuspended in complete medium (CM) (NBM, 0.01% GlutaMax, 0.04% B27 Supplement, 0.1% NT4, 0.01% Pen/Strep, 0.1% Glial cell-line Derived Neurotrophic Factor (GDNF)). 50 μl of the cell suspension was pipetted on each coverslip, and the coverslips were incubated for 30 min at 37°C. Finally, 2 ml of CM was added to each well. The culture has been incubated overnight in a 37°C incubator with 5% CO_2_. All phosphate-buffered saline solution (PBS) used during the culture preparation was depleted in Ca^2+^ and Mg^2+^. DRG neurons have been seeded on 13 mm coverslips, previously coated with poly-L-lysine and laminin. These experiments were approved by the KU Leuven Ethical Committee Laboratory Animals under project number (CMM-184/2021).

### Statistical analysis

Each experiment has been performed on at least 3 independent coverslips, with at least 100 total cells per condition. The statistical tests were performed using GraphPad Prism 10 software (chi-square test for proportions, Mann-Whitney test for independent skewed distributions). The definitive outliers have been identified and removed using the *Identify outliers* function of GraphPad Prism (ROUT method, Q = 0.1%). The number of Ca^2+^ transients per cell are represented as mean ± SEM, while the median amplitude of the transients is represented as box plots, where the middle line corresponds to the median value, the limits of the box correspond to the Q1 and Q3 quartiles, and the whiskers represent the 10% - 90% confidence intervals. In the figures, the asterisks indicate the level of statistically significant differences (*, p < 0.05 and **, p < 0.01).

### Reagents

The following solutions have been used throughout the measurements: LNP preparations (3.3 x 10^8^ – 10^11^ particles/mL), AITC 100 μM (Sigma-Aldrich), HC-030031 100 μM (Sigma-Aldrich), A-967079 10 μM (Tocris Bioscience), ruthenium red (RR) 10 μM (Sigma-Aldrich), pregnenolone sulfate 50 μM (Sigma-Aldrich), capsaicin 1 μM (Sigma-Aldrich), thapsigargin 2 μM (Tocris Bioscience), KCl 50 mM (Sigma-Aldrich). All measurements were performed in standard Krebs solution containing in mM: 150 NaCl, 6 KCl, 1.5 CaCl_2_ x 2H_2_O, 1 MgCl_2_ x 6H_2_O, 10 glucose, 5 HEPES buffer, 5 MES buffer (pH adjusted using NaOH). For the measurements performed in the absence of extracellular Ca^2+^, the Krebs solution contains the molar equivalent of CaCl_2_ in EGTA (pH adjusted using KCl).

## Acknowledgements

We thank Melissa Benoit for the technical support and the members of the LICR for helpful discussions. The CHO-mTRPA1 cell line was kindly provided by Ardem Patapoutian (The Scripps Research Institute, USA). The Python script for the detection of Ca^2+^ transients was developed in collaboration with Tamas Barath (KBC, Department of Advanced Data Analytics and Modelling, Belgium). This work was supported by grants from the FWO (G0AAL24N). GB and KT are partners of the FWO Scientific Research Network CaSign (W0.014.22N).

## Data availability statement

All data generated or analyzed during this study are included in the supporting files and online in the Research Data Repository of KU Leuven [89]. The Python script used to identify the Ca^2+^ transients is available on GitHub [86].

## Figures and tables

**Figure S1.**
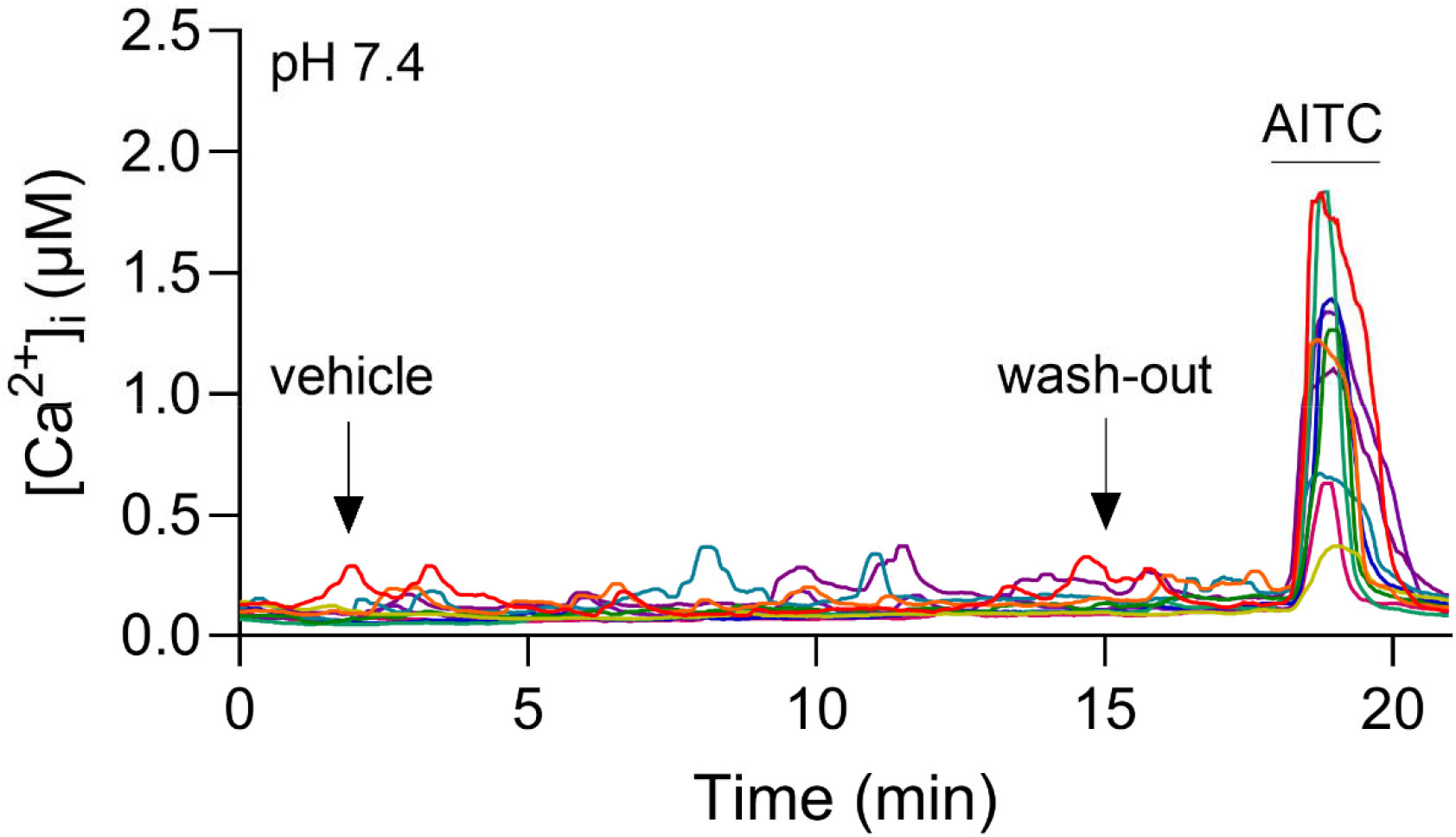
Representative traces of Ca^2+^ responses elicited by the injection of the vehicle solution (corresponding to LNP 10^11^ p/mL) in CHO-mTRPA1 cells at pH 7.4.

**Figure S2.**
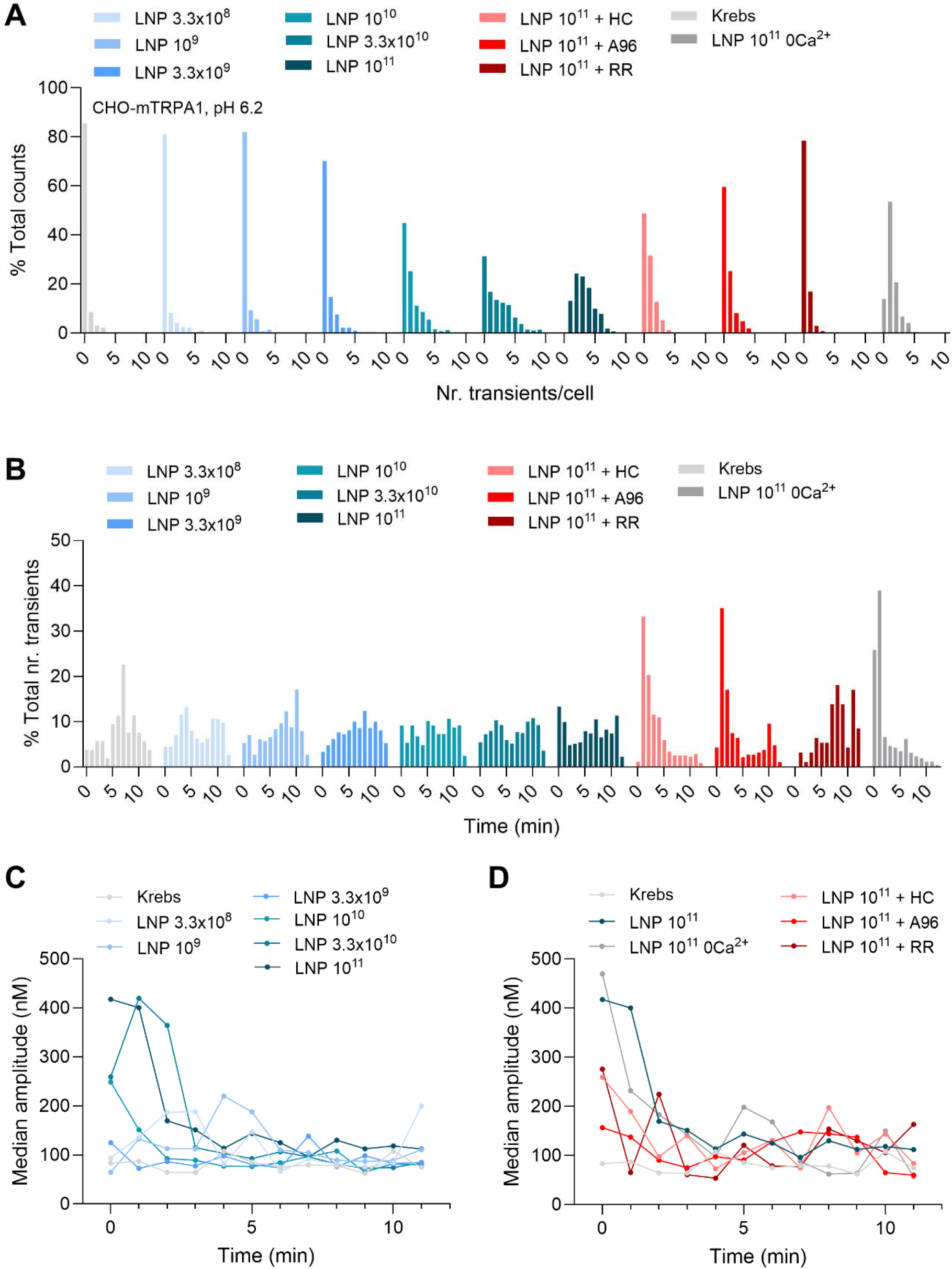
**A** Histogram of the relative distribution of the number of cells responding with a different number of Ca^2+^ transients per cell, identified in the whole population of responding CHO-mTRPA1 cells at pH 6.2 (binning of 1 Ca^2+^ transient per cell). **B** Histogram of the relative time distribution of the total number of Ca^2+^ transients identified throughout the application window in the whole population of responding CHO-mTRPA1 cells at pH 6.2 (binning of 1 min). **C** Median amplitude of Ca^2+^ responses measured for each minute of LNP application throughout the application window for different concentrations of LNPs in the whole population of responding CHO-mTRPA1 cells at pH 6.2. **D** Median amplitude of Ca^2+^ responses measured for each minute of LNP application throughout the application window for 10^11^ p/mL LNPs in the presence or absence of different TRPA1 channel blockers and in the absence of extracellular Ca^2+^, in the whole population of responding CHO-mTRPA1 cells at pH 6.2.

**Figure S3.**
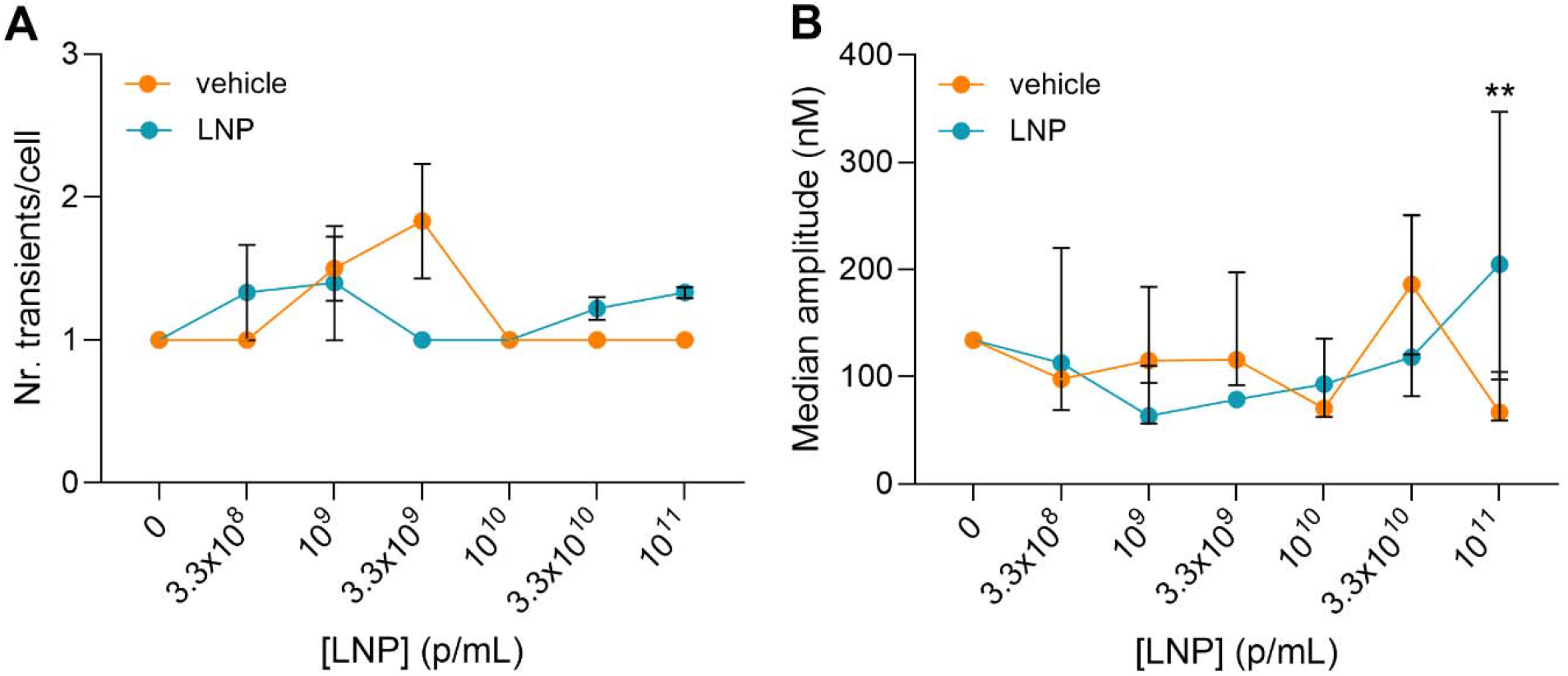
Dose response curves of the percentage of (A) number of Ca^2+^ transients and (B) median amplitude of Ca^2+^ transients in responding CHO-WT cells at pH 6.2. The asterisks correspond to the comparison between each concentration of LNP and the corresponding vehicle solution.

**Figure S4.**
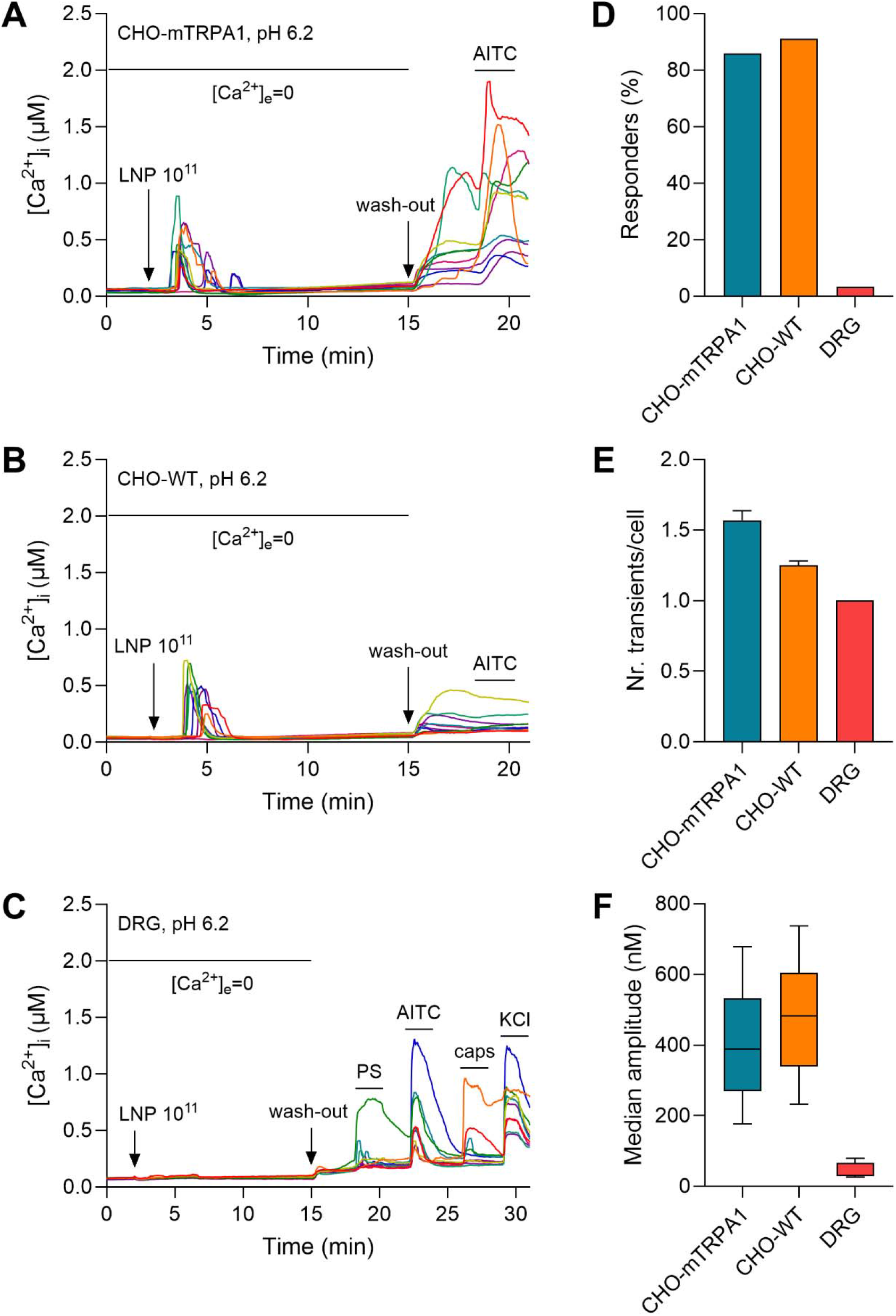
**A-C** Representative traces of Ca^2+^ responses induced by LNP 10^11^ p/mL in the absence of extracellular Ca^2+^ in (A) CHO-mTRPA1 cells, (B) CHO-WT cells, and (C) DRG neurons, at pH 6.2. **D-F** Quantification of Ca^2+^ responses induced by 10^11^ p/mL in the absence of extracellular Ca^2+^ in CHO-mTRPA1 cells, CHO-WT cells, and DRG neurons, at pH 6.2: (D) percentage of responding cells, (E) number of transients and (F) median amplitude of the Ca^2+^ transients induced in responding cells.

**Figure S5.**
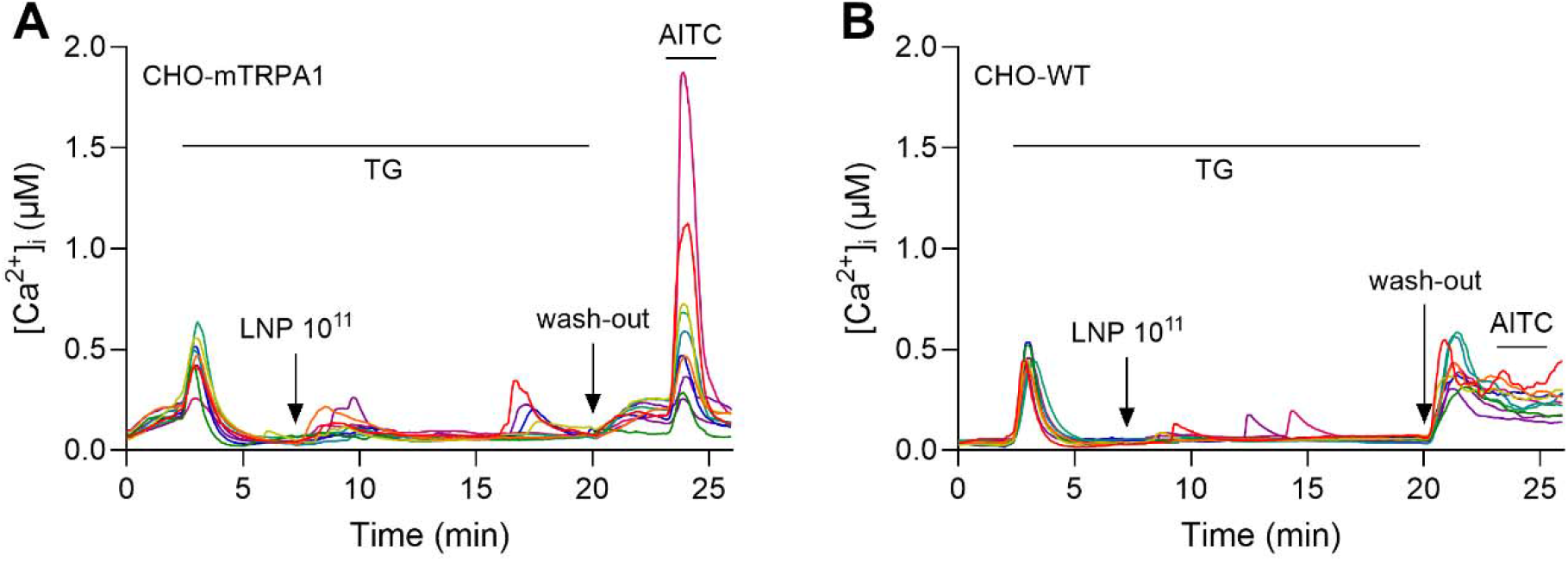
Representative traces of Ca^2+^ responses induced by LNP 10^11^ p/mL upon ER depletion with thapsigargin in (A) CHO-mTRPA1 cells and (B) CHO-WT cells, at pH 6.2.

